# A major update of the genome assembly of Indian Eri silkmoth, *Samia ricini*

**DOI:** 10.1101/2025.01.09.632291

**Authors:** Jung Lee, Mana Okamoto, Rin Kawagoe, Toru Shimada

**Affiliations:** Gakushuin University, Faculty of Science, Department of Life Science, Mejiro 1-5-1, Toshima-ku, Tokyo, 171-8588, Japan

## Abstract

Indian eri silkmoth, *Samia ricini*, is a wild silkmoth whose silk occupies a significant economic position. In addition to its importance as an economic animal, *S. ricini* is also useful as a model species of Saturniidae. National BioResource of Japan (NBRP) maintains a *S. ricini* strain brought to Japan during WWII via Taiwan. Since we have previously published a draft genome assembly of *S. ricini*, we have attempted to construct a chromosome-level genome assembly to facilitate genetic studies of *S. ricini*. We successfully constructed a chromosome-scale genome assembly by exploiting two long-read-based technologies, HiFi reads and optical genome mapping.

Furthermore, we performed functional annotations of the genome assembly, i.e., repeat annotation, transcriptome-based gene prediction, ATAC-seq, and PIWI-interacting RNA (piRNA)-targeted small RNA-seq. The assembly harbours 16,226 protein-coding genes and 636 piRNA clusters across three tissues: ovaries, testis, and embryos. ATAC-seq data comprehensively detected open chromosome regions, which will benefit when CRISPR/Cas9-mediated genome editing is conducted.

## Background & Summary

In India, many silkworm species other than *Bombyx mori* (mulberry silkworm) are reared for silk production. As far as we know, such species belong to the family Saturniidae. *Samia ricini*, eri silkworm, represents one example. Similar to *B. mori, S. ricini* is completely domesticated species and have no wild populations. Of the saturniid silkworms reared in India, *S. ricini* is the biggest cocoon producer^1^. In addition to that, there is a culture of eating the pupae of *S. ricini*, in the northern regions of India^2^. Since FAO (Food and Agriculture Organization of the United Nations) reported that insect diets are one of the practical solutions to the food problems^3^, there is growing interest in entomophagy. Therefore, *S. ricini* is expected to be in increasing demand not only as a sericultural insect but also as a food insect.

*S. ricini* also has another side to it as a model organism for the Saturniidae family. *S. ricini* is exceptionally multivoltine species in the family Saturniidae and can be reared throughout the year. Moreover, *S. ricini* is the only insect that genome editing has already been announced as being possible so far^4^. In view of this situation, the development of highly accurate whole-genome annotation information on *S. ricini* would be of great benefit to entomology.

We have published a draft genome sequence of *S. ricini* in 2020^5^, where 14 chromosomes of *S. ricini* were split into 155 contigs. We first attempted to improve the assembly accuracy without adding extra sequence data. We re-assembled the *S. ricini* genome using the continuous long read data and short read data from Lee et al., 2020^5^ (DRS137258^6^ and DRS137260^7^). *The following s*caffolding was conducted with optical genome mapping data (Table 1)^8^. In the resulting assembly (Sr_NBRP_v2.0; GCA_014132275.2)^9^, the number of sequences and chromosomes were equal, and we were able to construct a ‘chromosome level’ assembly for the time being. However, the BUSCO completeness was low (Table 2), suggesting that there were many undetermined regions. In addition, the total gap size was approximately 4.5 Mbp, which seemed to be the performance limit of Noisy Long reads. Therefore, we obtained approximately 30 Gbase (coverage x60) of data using HiFi reads and performed assembly and scaffolding. As a result, we have succeeded in constructing a chromosome-scale assembly containing only 400 bp gaps (Table 1; Sr_NBRP_v3.0; AP038896–AP038909^10–23^).

**Table 1.** Basic statistics of the genome assemblies of *S. ricini*.

| Assembly | Accession | Strain | Level | No. of sequences | Total bases | N50 [bp] | Gap [bp] | BUSCO completeness [%] | Source |
| --- | --- | --- | --- | --- | --- | --- | --- | --- | --- |
| Sr_NBRP_v1.0 | GCA_014132275.1 | White | Contig | 155 | 450,479,495 | 21,366,385 | 0 | 97.6 | Lee et al., 2020 |
| Sr_NBRP_v2.0 | GCA_014132275.2 | White | Chromosome | 14 | 452,483,983 | 33,135,288 | 4,566,028 | 96.4 | This Study |
| Sr_NBRP_v3.0 | AP038896-AP038909 | White | Chromosome | 14 | 453,650,863 | 33,371,992 | 400 | 98.8 | This Study |

**Table 2.** Sequence read information used in this study. Since sequence runs of IDs 19, 23–27 were performed on long read sequencer, ‘Length’ and ‘Read pairs after trimming’ cells are empty.

| ID | Run accession | biosample accession | Purpose | No. of spots | Length | Read pairs after trimming | tissue | age | Source |
| --- | --- | --- | --- | --- | --- | --- | --- | --- | --- |
| 1 | DRR596219 | SAMD00819509 | RNA-seq | 49,533,284 | 101PE | 23,747,007 | anterior and middle silk gland | 5th instar 3rd day larvae | This study |
| 2 | DRR596220 | SAMD00819510 | RNA-seq | 62,957,650 | 101PE | 30,706,997 | epidermis | 3rd instar 0th day larvae | This study |
| 3 | DRR596221 | SAMD00819511 | RNA-seq | 57,939,494 | 101PE | 28,209,875 | epidermis | 4th instar 1st day larvae | This study |
| 4 | DRR596222 | SAMD00819512 | RNA-seq | 45,644,604 | 101PE | 22,195,994 | epidermis | 5th instar 3rd day larvae | This study |
| 5 | DRR596223 | SAMD00819513 | RNA-seq | 47,654,200 | 101PE | 23,060,565 | midgut | 5th instar 3rd day larvae | This study |
| 6 | DRR596224 | SAMD00819514 | RNA-seq | 57,351,070 | 101PE | 27,645,513 | posterior silk gland | 5th instar 3rd day larvae | This study |
| 7 | DRR596225 | SAMD00819515 | RNA-seq | 15,522,314 | 150PE | 15,413,240 | hemocytes | 5th instar larvae | This study |
| 8 | DRR596226 | SAMD00819516 | RNA-seq | 15,351,834 | 150PE | 15,260,640 | fat body | 5th instar larvae | This study |
| 9 | DRR596227 | SAMD00819517 | RNA-seq | 16,354,434 | 150PE | 16,260,987 | gonads | 5th instar larvae | This study |
| 10 | DRR596228 | SAMD00819518 | RNA-seq | 14,152,184 | 150PE | 14,062,249 | malpighian tubules | 5th instar larvae | This study |
| 11 | DRR596229 | SAMD00819519 | RNA-seq | 13,758,095 | 150PE | 13,673,816 | midgut | 5th instar larvae | This study |
| 12 | DRR596230 | SAMD00819520 | RNA-seq | 14,007,357 | 150PE | 13,904,310 | adult's 9th ganglion | immediately after eclosion | This study |
| 13 | DRR596231 | SAMD00819521 | RNA-seq | 14,743,955 | 150PE | 14,639,299 | adult's 9th ganglion | after copulation | This study |
| 14 | DRR596232 | SAMD00819522 | RNA-seq | 14,351,152 | 150PE | 14,267,229 | nerve chord | 5th instar larvae | This study |
| 15 | DRR596233 | SAMD00819523 | RNA-seq | 15,453,946 | 150PE | 15,184,662 | adult's brain | immediately after eclosion | This study |
| 16 | DRR596234 | SAMD00819524 | RNA-seq | 15,236,362 | 150PE | 14,781,588 | adult's brain | after copulation | This study |
| 17 | DRR596235 | SAMD00819525 | RNA-seq | 14,459,033 | 150PE | 14,243,379 | midgut | 5th instar larvae | This study |
| 18 | DRR596236 | SAMD00819526 | RNA-seq | 14,520,390 | 150PE | 14,379,533 | midgut | 5th instar larvae | This study |
| 19 | DRR396182 | SAMD00515715 | iso-seq | 9,094,853 | - | - | early embryo | 48 hours post oviposition | This study |
| 20 | DRR396184 | SAMD00515715 | small RNA-seq | 48,219,633 | 50PE | 48,219,633 | early embryo | 48 hours post oviposition | This study |
| 21 | DRR396185 | SAMD00515716 | small RNA-seq | 25,347,659 | 50PE | 25,347,659 | Pupal ovary | 7 days post pupation | This study |
| 22 | DRR396186 | SAMD00515717 | small RNA-seq | 29,625,406 | 50PE | 29,625,406 | Pupal testis | 7 days post pupation | This study |
| 23 | DRR596218 | SAMD00819399 | Genomic DNA-seq | 1,776,772 | - | - | whole body | 5th instar larvae | This study |
| 24 | DRR213158 | SAMD00207788 | Genomic DNA-seq | 777,113 | - | - | silk gland | 5th instar larvae | Lee et al., 2020 |
| 25 | DRR213159 | SAMD00207788 | Genomic DNA-seq | 706,515 | - | - | silk gland | 5th instar larvae | Lee et al., 2020 |
| 26 | DRR213156 | SAMD00207788 | Genomic DNA-seq | 925,105 | - | - | silk gland | 5th instar larvae | Lee et al., 2020 |
| 27 | DRR213157 | SAMD00207788 | Genomic DNA-seq | 853,458 | - | - | silk gland | 5th instar larvae | Lee et al., 2020 |
| 28 | DRR213150 | SAMD00207786 | Genomic DNA-seq | 36,348,660 | 126PE | 35,191,810 | silk gland | 5th instar larvae | Lee et al., 2020 |

Transcriptome-based gene prediction and functional annotation were also performed, identifying 16,226 protein-coding genes in this assembly. For the future application of the CRISPR/Cas9, we performed embryonic ATAC-seq to associate the positions of protein-coding genes and open chromatin regions since the enzymatic activity of Cas9 is less efficient in heterochromatin regions^24^. In addition to ATAC-seq, we performed piRNA-targeted small RNA-seq to identify piRNA clusters (piCs) since piRNA is involved in the early development of Lepidoptera^25^.

## Methods

### Insects

*S. ricini* (White strain) was provided from NBRP silkworm (https://shigen.nig.ac.jp/silkwormbase/). The *S. ricini* larvae were fed on fresh castor leaves under a long-day condition (16 h light / 8 h dark) at 25°C.

### Genome assembly of Sr_NBRP_v2.0 (GCA_014132275.2)^9^

PacBio continuous long read (CLR) data (DRS137260)^7^ were subjected to Flye (v 2.9)^26^ with default setting. After the draft assembly step, raw CLR data was reused for polishing with Racon^27^. In this case, consensus calling by Racon was converged by one attempt. The resulting polished draft sequences were subjected to Pilon (v 1.22) 28 for further polishing. For polishing with Pilon, 2 paired-end libraries (DRR213145 and DRR244931)^29^ and 9 mate-pair libraries (DRR213147, DRR213148, DRR213149, DRR213150, DRR213151, DRR213152, DRR213153, DRR213154, and DRR213155)^29^ were used. The polished assembly was scaffolded with optical genome mapping as previously described^30^: Genomic

DNA was isolated from the pupa immediately after the pupation for the optical genome mapping. DNA isolation was conducted using the Bionano Prep Animal Tissue DNA Isolation Fibrous Tissue Protocol (Bionano Genomics). We used two enzymes, namely, DLE-1 and Nt.BsqQI for labelling. The labelling procedure was conducted according to the Bionano Prep Direct Label and Stain Protocol. The labelled samples were scanned on the Bionano Saphyr system using Saphyr Chip G2.3. The obtained data were analyzed using Bionano Access (v 1.8.2)^31^ and Bionano Solve (v 3.8.2)^32^. In the two-enzyme pipeline, the process of creating the .cmap files was the same as in the one-enzyme pipeline. The resulting two .cmap files are submitted to runTGH.R (also bundled with Bionano Solve) with default settings. The .cmap files were deposited in DDBJ. The basic statistics of this assembly are summarised in Table 1.

### Genome assembly of Sr_NBRP_v3.0 (AP038896–AP038909)^10–23^

Genomic DNA was extracted from whole body of a male final instar larvae of S. ricini using NucleoBond HMW DNA (MACHEREY-NAGEL). A SMRTbell library was prepared with SMRTbell library preparation 3.0. HiFi reads were obtained on the Revio platform. The HiFi reads were assembled with Hifiasm (v 0.16.1-r375)^33^. The resulting draft assembly was scaffolded with the above-mentioned optical genome mapping data. To close the gaps, which were bridged by optical genome mapping reads, we attempted to use error-corrected CLR (DRS137260)^7^ :the raw CLRs were subjected into the “correct” module in PECAT (v 0.0.3)^33^ with default settings. The gaps in the scaffolded assembly were patched to the error-corrected CLRs using “GapFiller” module in QuarTeT (v 1.2.0)^34^. To close remaining gaps, we patched the previous assembly (Sr_NBRP_v2.0; GCA_014132275.2)^9^ using “AssemblyMapper” module in QuarTeT (v 1.2.0)^34^. Finally, we polished the resulting assembly with HiFi reads with NextPolish2 (v 0.2.1)^35^ with default settings. Winnowmap (v 2.03)^36^ was used with default parameters to align HiFi reads to the temporary genome assembly after “AssemblyMapper.” Yak (v 0.1-r69)^33^ was used with default parameters to create k-mer hash tables from short read data (DRR213150)^29^ which were mandatory inputs for NextPolish2. Schematic diagram of assembly process was summarised in Fig. 1. The assembly gaps were visualised by chromoMap (v 4.1.1)^37^ in Fig. 2a. All sequence reads data used in this study were summarised in Table 2.

**Fig 1.**
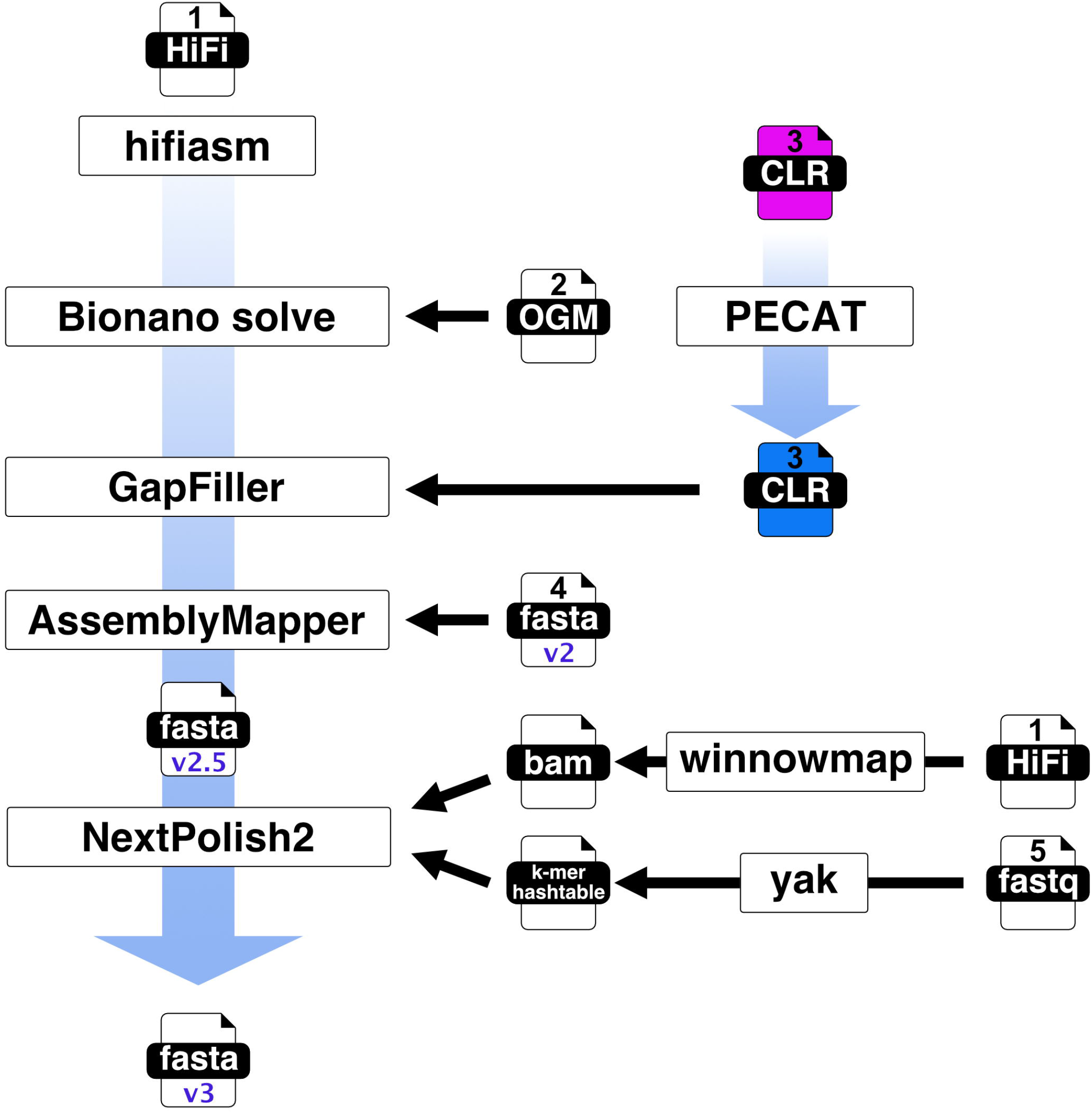
Genome assembly pipeline used in this study. The numbers 1 to 5 in the figure identify one of the five files used as input (1, HiFi reads; 2, Optical genome mapping data; 3, continuous long reads; 4, previous version of genome assembly; 5, short reads).

**Fig 2.**
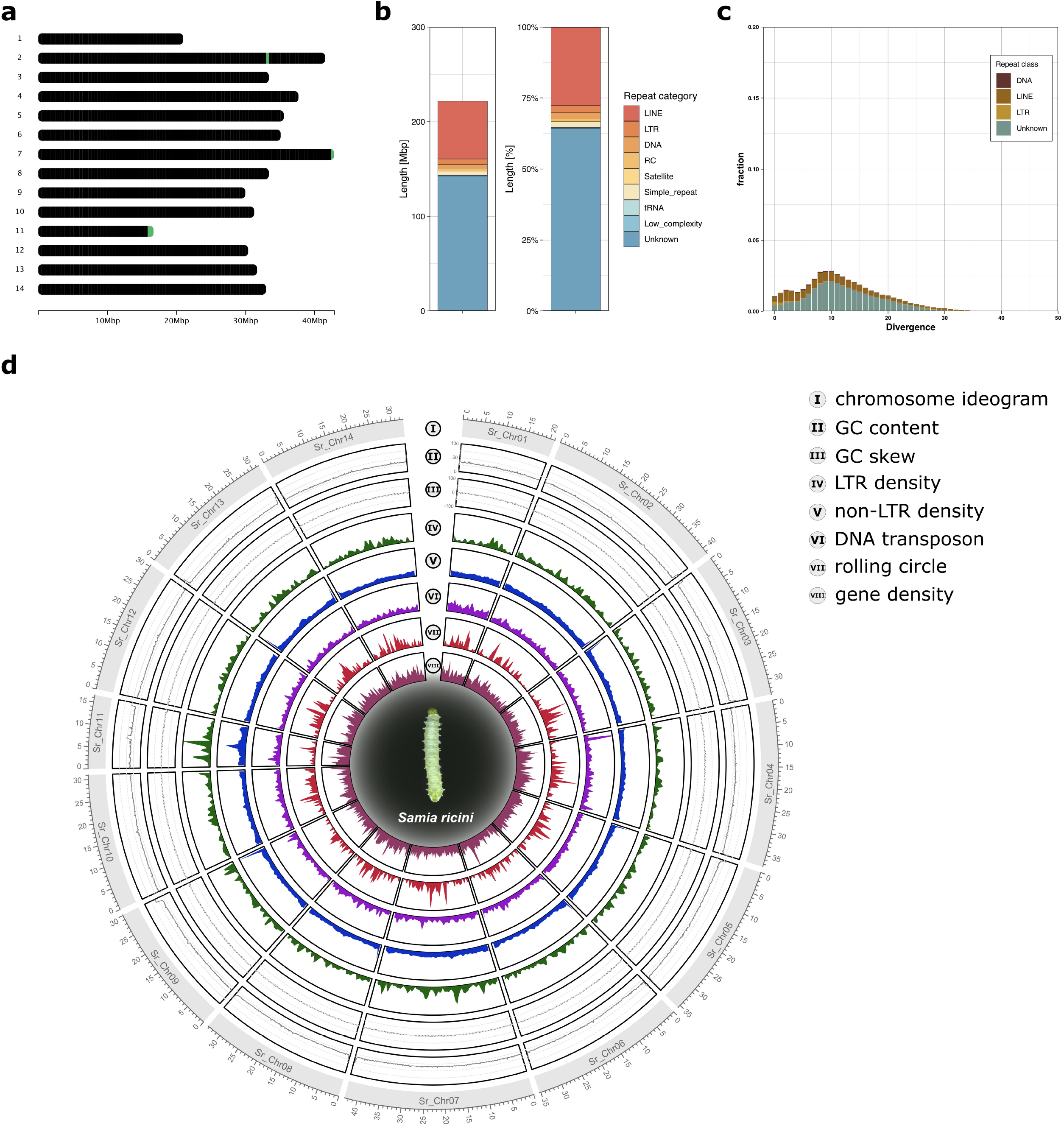
A chromosome-scale genome assembly of S. ricini. (a)Gap distribution on the genome assembly of *S. ricini*. There is one gap each on both chromosome 2 and 7. There are two gaps in close proximity in the terminal region of chromosome 11. (b)Repeat contents of the genome assembly. Amount (Mbp; left) and fractions (right) by repeat category were shown. (c)Repeat landscape graph based on divergence data of repetitive elements. (d)Summary of *S. ricini* genome characteristics. The outermost to the innermost circle are I. chromosome ideograms; II. GC content; III. GC skew; IV. LTR element density; V. non-LTR retrotransposon density; VI. DNA transposon density; VII. rolling circle density; and VIII. gene model density.

### Repetitive elements annotation in the genomes

Repetitive annotation of the *S. ricini* genome was briefly summarised here: repetitive elements in the genome assembly were identified using RepeatModeler (v 2.0.4)^38^ with the “-LTRstruct” option for performing an LTR structural search. The annotated elements were masked using RepeatMasker (v 4.1.2)^39^ with ‘-lib’ option to specify “consensi.fa.classified” file generated by RepeatModeler. The amount of annotated repetitive elements in the assembly was summarised in Fig. 2b. We realised that absence of SINE in the genome can characterise the repetitive elements of *S. ricini* genome (Fig. 2b). Repeat Landscape graph using the divergence data was also generated (Fig. 2c). Among the annotated repetitive elements, LTR, non-LTR (LINE or SINE), DNA transposons, and rolling circles were extracted and the density information of those repetitive groups is visualized by circlize (v 0.4.16)^40^ (Fig. 2d).

### RNA sample preparation for RNA-seq, small RNA-seq, and Iso-seq

All RNA samples were prepared precisely as previously described^30^. Total RNA was extracted using TRIzol reagent (Invitrogen) according to the manufacturer’s protocol. Embryos were sampled 48 hours after oviposition. Testis and ovary-derived RNA samples were subjected to RNA-seq and small RNA-seq (sRNA-seq), respectively. Embryonic RNA samples were subjected to sRNA-seq and Iso-seq respectively.

### Library preparation for RNA-seq, sRNA-seq and Iso-seq

The sRNA-seq library was prepared using TruSeq small RNA kit (illumina) according to the manufacturer’s protocol with a slight modification. To target piRNA, a region of 147-158 nucleotides was extracted in the purification step of the cDNA construct using BluePippin (Sage Science). The constructed library was sequenced on the illumina HiSeq 2500 platform (illumina). Except for an RNA sample from embryos, the RNA-seq libraries were prepared using TruSeq stranded mRNA kit (illumina) according to the manufacturer’s protocol. The embryonic RNA-seq library was prepared using NEBNext Poly(A) mRNA Magnetic Isolation Module (New England BioLabs) and the NEBNext^®^ Ultra™ ll Directional RNA Library Prep Kit (New England BioLabs) according to the manufacturer’s protocol. The constructed RNA-seq libraries were sequenced on the illumina NovaSeq6000 platform (illumina). For Iso-seq, the library was constructed using Sequel Iso-seq Express Template Prep (Pacific Bioscience) according to the manufacturer’s protocol. The constructed library was sequenced on the PacBio Sequel platform (PacBio).

### Transcriptome-based gene prediction

BRAKER3 (v 3.0.8) was used for gene prediction^41,42^. The RNA-seq and Iso-seq data were submitted to BRAKER3 separately^43^, and Tsebra finally merged their respective prediction^44^. The detailed information on transcriptome data is summarised in Supplementary Table 2. Quality trimming for short read data was conducted using fastp (v 0.20.1)^45^ with following options: ‘-q 28 -l 80. ‘ Trimmed short read data were submitted to BRAKER3 using the ‘--rnaseq_sets_ids’ option. The short reads were aligned to the genome assembly by hisat2 (v 2.2.1)^46^. Iso-seq data were generated consensus for each read cluster according to the following procedure47: Iso-seq subreads were converted to circular consensus sequences (ccs) using ccs v 6.4.0 with options ‘--minLength 10 --maxLength 100000 -- minPasses 0 --minSnr 2.5 --minPredictedAccuracy 0.0.’ lima (v 2.7.1) was used to remove primer sequences from the CCSs with options ‘--isoseq --peek-guess --ignore-biosamples.’ After the trimming of adaptors, PolyA tail trimming and concatemer removal were performed by isoseq3 (v 3.8.2) in ‘refine’ mode with option ‘--require-polya.’ Finally, isoform-level clustering was conducted by isoseq3 in ‘cluster’ mode with option ‘--use-qvs.’ The resulting clustered.bam file was submitted to BRAKER3. The basic metrics of gene models were summarised in Table 3. The resulting gene models were also submitted to BUSCO analysis^30^, scoring 98.4% completeness (Table 4). BUSCO analysis on the genome assembly was also conducted to compare completeness scores of both gene models and genome assembly^30^ (Table 4).

**Table 3.** Basic statistics of the predicted protein-coding genes.

|  |  |
| --- | --- |
| No. of protein coding gene | 16,226 |
| Average CDS length [bp] | 1512.4 |
| Average exon length [bp] | 228.003 |
| Average intron length [bp] | 1822.58 |

**Table 4.** BUSCO completeness scores of the predicted protein-coding genes and the genome assembly.

|  | Complete (Single) | Complete (Duplicated) | Fragmented | Missing |
| --- | --- | --- | --- | --- |
| Gene model | 5137 (97.2%) | 65 (1.2%) | 13 (0.2%) | 71 (1.4%) |
| Genome assembly | 5204 (98.4%) | 13 (0.2%) | 14 (0.3%) | 55 (1.1%) |

### Functional annotation of gene models

The deduced amino acid sequences of gene models were submitted to EnTAP6 for functional annotation. A protein similarity search was conducted against the latest complete UniProtKB/TrEMBL protein data set and complete UniProtKB/Swiss-Prot data set using diamond (v 0.9.14)^48^. A protein orthology search was also conducted against the EggNOG databases^30^ to assign Gene Ontology (GO), KEGG terms, and protein domains from pfam^49^ and smart^50^. Additional family and domain search was performed against TIGRFAM^51^, SFLD^52^, HAMAP^53^, CDD^54^, SUPERFAMILY^55^, PRINTS^56^, PANTHER^57^, and Gene3d^58^ using InterProScan (v 5.70-102)^59^. The results of functional annotation are summarized in Table 5. The top 10 GOs assigned to the gene models are shown in Fig. 3a without distinguishing between molecular function, biological process, and cellular component. The top 10 GOs for each category are shown in Fig. 3b–d.

**Table 5.** Brief summary of functional annotation. ^*^When the query sequences were aligned to sequences whose description contains any of conserved/predicted/unnamed/hypothetical/putative/unidentified/uncharacterized/unknown/uncultured/uninformative, such alignment was categorized as “uninformative,” and the query sequence was treated as an unannotated sequence. ^**^In this column, queries with at least one GO term were treated as “Informative,” while queries without GO terms were treated as “Uninformative.” “Unaligned” in this column means queries without protein family assignment. ^***^”Annotated” means at least one match yielded from any of the databases. “Unannotated” means no match yielded from all databases.

|  |  | Similarity search |  |  | Ontology search |  |  |  |
| --- | --- | --- | --- | --- | --- | --- | --- | --- |
|  |  | EggNOG | TrEMBL | Swiss-prot | EggNOG** | InterPro | total |  |
| aligned | informative | 12843 | 12003 | 7872 | 10886 | 12,906 | annotated*** | 14430 |
|  | uninformative* | 0 | 2101 | 182 | 3046 | - |  |  |
| unaligned |  | 3383 | 2122 | 8172 | 2294 | 3,320 | unannotated*** | 1796 |

**Fig 3.**
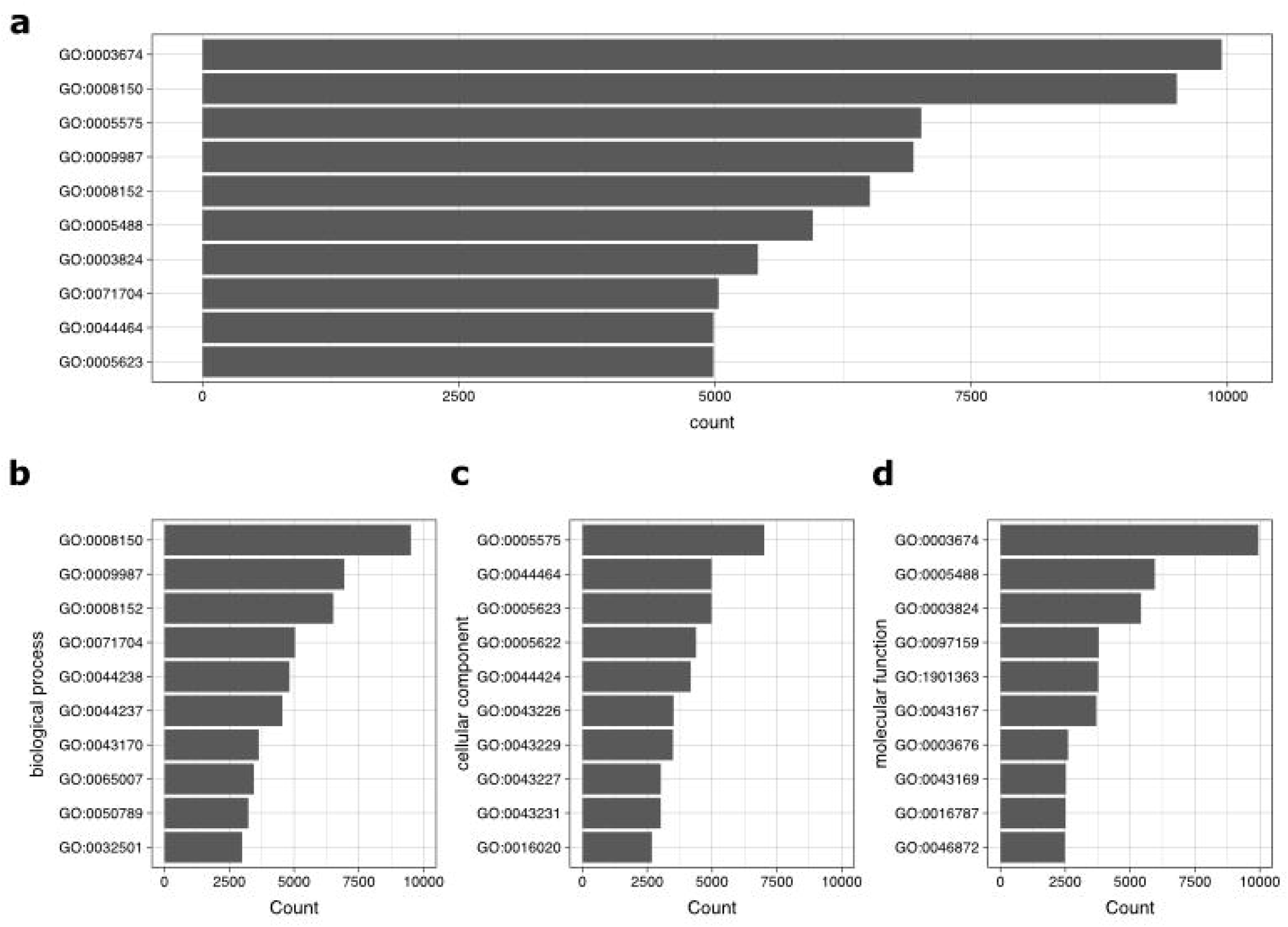
The top 10 GO assignments of gene models. (a) Overall top 10 GO assignments to gene models (b)Top 10 “biological process” GO assignments to gene models (c)Top 10 “cellular component” GO assignments to gene models (d) Top 10 “molecular function” GO assignments to gene models

### ATAC library preparation and data processing

Another batch of early embryo samples was subjected to Iso-seq, and sRNA-seq was subjected to ATAC-seq. Fragmentation and amplification of the ATAC-seq libraries were conducted according to Buenrostro et al. (2015)^60^. The constructed libraries were sequenced on the Illumina HiSeq. ATAC-seq reads were pretreated with fastp and mapped to the genome with bwa-mem2 (v 2.2.1)^61^. Alignments containing mismatches were then removed using bamutils (v 0.5.9)^62^. Next, we removed duplicated reads using GATK MarkDuplicates (v 4.1.7)^63^. The resulting bam files were converted to bigwig files using deepTools bamCoverage (v 3.5.1)^64^. Heatmap was created using deepTools computeMatrix, and the starting point of the gene model was set to the reference point (Fig. 4).

**Fig 4.**
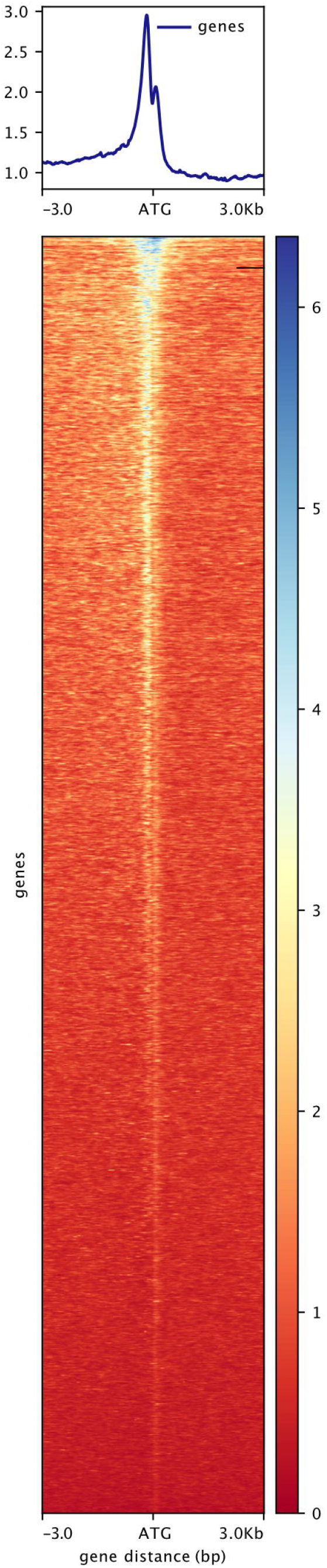
Heatmap around gene bodies of ATAC-seq on early embryos.

### Small RNA mapping

The small RNA reads were trimmed using Trim Galore (v 0.6.6)^65^ in small RNA mode. The trimmed small RNA reads were mapped to the assembled transcriptome, allowing up to 3 nucleotide mismatches using Hisat2 (v 2.1.0)^46^ and ngsutils (v 0.5.9)^62^. The information for each library is summarised in Table 2.

### piRNA cluster detection

The piC detection was performed as previously described^30,66^. proTRAC (v 2.4.4)^66^ was used with options ‘-clsize 5000 -pimin 23 -pimax 29 -1Tor10A 0.3 -1Tand10A 0.3 -clstrand 0.0 -clsplit 1.0 -distr 1.0-99.0 -spike 90-1000 -nomotif -pdens 0.05.’ As a result, we successfully identified 560 piRNA clusters in the three tissues (Fig. 5). The identity of piC is defined by the two nearest gene models. If multiple piCs were predicted between such two genes, such piCs were treated as a single piC. The genomic positions of piCs identified in testes, ovaries, and early embryos were visualized by RIdeogram (v 0.2.2)^67^ (Fig. 5a). The aggregation relationship of those piCs was visualized by ComplexUpset (v 1.3.3)^68^ (Fig. 5b).

**Fig 5.**
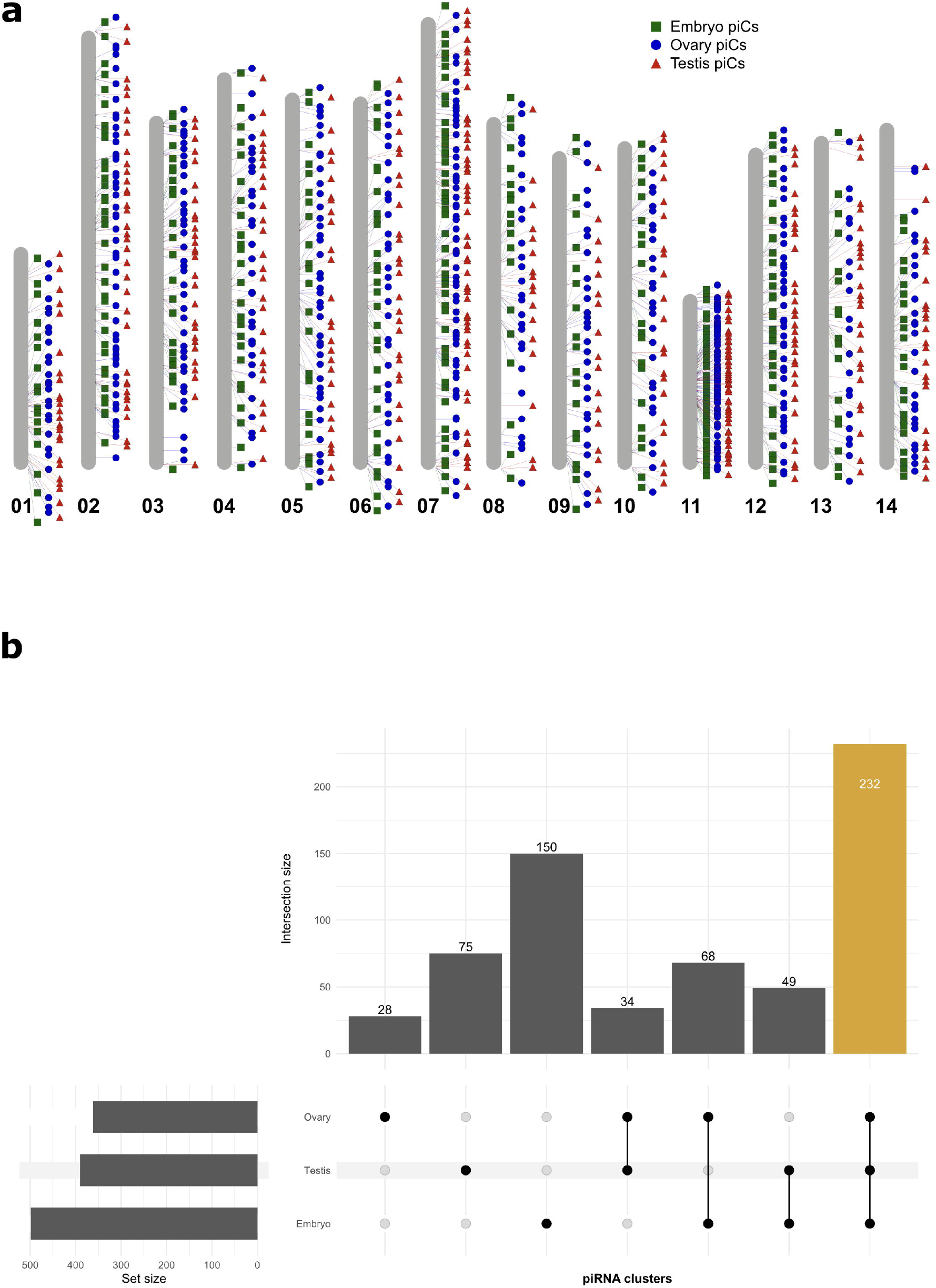
piRNA clusters on *S. ricini* genome. (a)piCs distribution detected in early embryos (box), pupal ovary (circle), and pupal testis (triangle) (b)UpSet plot visualising piCs that are each assigned to each tissue. The vertical bars correspond to the intersections. When the circles corresponding to tissues are connected by a line, the bar above circles represents the number of piCs commonly identified in concerning tissues. For example, the yellow bar indicates the number of piCs identified in all tissues. The identity of piCs was defined by the nearest two gene models: When comparing piCs identified in different tissues, if the nearest upstream and downstream gene models are the same, those piCs were treated as the same piC.

## Data Records

The raw sequence data reported in this paper have been deposited in DDBJ. Genomic data for the draft assembly were deposited under the accession code DRP012461^69^. Except for embryonic Iso-seq data, all transcriptome data were also registered under the accession code DRP012461. Embryonic Iso-seq data, small RNA-seq data and ATAC-seq data are available under the accession code DRP009926 [DRR396182, DRR396184, DRR396185, and DRR396186]^70^. Annotated gene models and piRNA distribution maps have been deposited to the figshare repository^71^.

## Technical Validation

BUSCO (v 5.4.6)^72^ with lepidoptera_odb10 lineage dataset was used to assess the quality of gene models. For comparison, the results are summarised in Table 4, together with the results of BUSCO analysis for the genome assembly. 98.4% of the complete and single-copy BUSCO sequences were present in the gene models, while 98.6% of the complete and single-copy BUSCO sequences were in the genome assembly. BUSCO completeness scores were almost the same between the genome assembly and the gene model, suggesting that the gene prediction process is highly accurate across all genome regions.

## Code Availability

Programs exploited in this study were executed with the default parameters except where otherwise specified in the Methods section. No custom code was used during this study.

## Acknowledgments

Insects were donated from Kyushu University and Shinshu University according to a Grant-in Aid “National BioResource Project (NBRP, RR2002), Silkworm Genetic Resources” for Scientific Research from the Ministry of Education, Science, Sports and Culture of Japan. This study was funded by Ellie Inc. This study was also supported by JSPS KAKENHI Grant Numbers 20K15535 and 24K17900 to J.L and JSPS KAKENHI Grant Number J18H03949 to T.S.

## Author contributions

J.L. designed the research plan, performed RNA extraction, analyzed the obtained data, and wrote the manuscript. T.S. also designed this research plan and performed the data analysis. M.O. and R.K. reared the larvae of *S. ricini* and prepared RNA samples. T.A.

## References

1. Central Silk Board. SERICULTURAL STATISTICS IN INDIA. https://texmin.nic.in/sites/default/files/Sericulture%20Statistics.pdf (2023).

2. Shrivastava, S. K. & Prakash, A. INSECTS FOR FOOD: AN UNEXPLORED BIORESOURCE FOR MAKE IN INDIA. J. Appl. Zool. Res vol. 27 https://cabidigitallibrary.org (2016).

3. Huis, A. et al. EDIBLE INSECTS Future Prospects Fo Food and Feed Security. vol. 171 (2013).

4. Lee, J., Kiuchi, T., Kawamoto, M., Shimada, T. & Katsuma, S. Accumulation of uric acid in the epidermis forms the white integument of Samia ricini larvae. PLoS One 13, e0205758 (2018).

5. Lee, J. et al. The genome sequence of Samia ricini, a new model species of lepidopteran insect. Mol Ecol Resour 21, 327–339 (2021).

6. Lee, J. NCBI Sequence Read Archive. http://identifiers.org/ncbi/insdc.sra:DRS137258 http://identifiers.org/ncbi/insdc.sra:DRS137258 (2020).

7. Lee, J. NCBI Sequence Read Archive. https://identifiers.org/ncbi/insdc.sra:DRS137260 https://identifiers.org/ncbi/insdc.sra:DRS137260 (2020).

8. Lee, J. et al. The genome sequence of Samia ricini, a new model species of lepidopteran insect. Mol Ecol Resour 21, 327–339 (2020).

9. Lee, J. & Shimada, T. NCBI GenBank. http://identifiers.org/ncbi/insdc.gca:GCA_014132275.2 (2023).

10. Lee, J. & Shimada, T. NCBI nucleotide. https://identifiers.org/ncbi/insdc:AP038896 (2025).

11. Lee, J. & Shimada, T. NCBI nucleotide. https://identifiers.org/ncbi/insdc:AP038897 (2025).

12. Lee, J. & Shimada, T. NCBI nucleotide. https://identifiers.org/ncbi/insdc:AP038898 (2025).

13. Lee, J. & Shimada, T. NCBI nucleotide. https://identifiers.org/ncbi/insdc:AP038899 (2025).

14. Lee, J. & Shimada, T. NCBI nucleotide. https://identifiers.org/ncbi/insdc:AP038900 (2025).

15. Lee, J. & Shimada, T. NCBI nucleotide. https://identifiers.org/ncbi/insdc:AP038901 (2025).

16. Lee, J. & Shimada, T. NCBI nucleotide. https://identifiers.org/ncbi/insdc:AP038902 (2025).

17. Lee, J. & Shimada, T. NCBI nucleotide. https://identifiers.org/ncbi/insdc:AP038903 (2025).

18. Lee, J. & Shimada, T. NCBI nucleotide. https://identifiers.org/ncbi/insdc:AP038904 (2025).

19. Lee, J. & Shimada, T. NCBI Nucleotide. https://identifiers.org/ncbi/insdc:AP038905 (2025).

20. Lee, J. & Shimada, T. NCBI Nucleotide. https://identifiers.org/ncbi/insdc:AP038906 (2025).

21. Lee, J. & Shimada, T. NCBI Nucleotide. https://identifiers.org/ncbi/insdc:AP038907 (2025).

22. Lee, J. & Shimada, T. NCBI Nucleotide. https://identifiers.org/ncbi/insdc:AP038908 (2025).

23. Lee, J. & Shimada, T. NCBI Nucleotide. https://identifiers.org/ncbi/insdc:AP038909 (2025).

24. Jain, S. et al. TALEN outperforms Cas9 in editing heterochromatin target sites. Nat Commun 12, 4–13 (2021).

25. Kiuchi, T. et al. A single female-specific piRNA is the primary determiner of sex in the silkworm. Nature 509, 633–636 (2014).

26. Kolmogorov, M., Yuan, J., Lin, Y. & Pevzner, P. A. Assembly of long, error-prone reads using repeat graphs. Nat Biotechnol 37, 540–546 (2019).

27. Vaser, R., Sović, I., Nagarajan, N. & Šikić, M. Fast and accurate de novo genome assembly from long uncorrected reads. Genome Res 27, 737–746 (2017).

28. Walker, B. J. et al. Pilon: An integrated tool for comprehensive microbial variant detection and genome assembly improvement. PLoS One 9, (2014).

29. Lee, J. NCBI Sequence Read Archive. https://identifiers.org/ncbi/insdc.sra:DRS137258 (2020).

30. Lee, J. et al. W chromosome sequences of two bombycid moths provide an insight into the origin of Fem. Mol Ecol 33, 1–12 (2024).

31. Bionano Genomics, I. Bionano Access Software. Preprint at https://bionano.com/access-software/.

32. Bionano Genomics, I. Bionano Solve software. Preprint at https://bionano.com/software-downloads/.

33. Cheng, H., Concepcion, G. T., Feng, X., Zhang, H. & Li, H. Haplotype-resolved de novo assembly using phased assembly graphs with hifiasm. Nat Methods 18, 170–175 (2021).

34. Lin, Y. et al. QuarTeT: A telomere-To-Telomere toolkit for gap-free genome assembly and centromeric repeat identification. Hortic Res 10, (2023).

35. Hu, J. et al. NextPolish2: A Repeat-aware Polishing Tool for Genomes Assembled Using HiFi Long Reads. Genomics Proteomics Bioinformatics 22, (2024).

36. Jain, C., Rhie, A., Hansen, N. F., Koren, S. & Phillippy, A. M. Long-read mapping to repetitive reference sequences using Winnowmap2. Nat Methods 19, 705–710 (2022).

37. Anand, L. & Rodriguez Lopez, C. M. ChromoMap: an R package for interactive visualization of multi-omics data and annotation of chromosomes. BMC Bioinformatics 23, (2022).

38. Flynn, J. M. et al. RepeatModeler2 for automated genomic discovery of transposable element families. Proc Natl Acad Sci U S A 117, 9451–9457 (2020).

39. Smit, A., Hubley, R. & Green, P. RepeatMasker Open-4.0. 2013–2015.

40. Gu, Z., Gu, L., Eils, R., Schlesner, M. & Brors, B. Circlize implements and enhances circular visualization in R. Bioinformatics 30, 2811–2812 (2014).

41. Stanke, M., Schöffmann, O., Morgenstern, B. & Waack, S. Gene prediction in eukaryotes with a generalized hidden Markov model that uses hints from external sources. BMC Bioinformatics 7, 62 (2006).

42. Stanke, M., Diekhans, M., Baertsch, R. & Haussler, D. Using native and syntenically mapped cDNA alignments to improve de novo gene finding. Bioinformatics 24, 637– 644 (2008).

43. Lomsadze, A., Burns, P. D. & Borodovsky, M. Integration of mapped RNA-Seq reads into automatic training of eukaryotic gene finding algorithm. Nucleic Acids Res 42, 1–8 (2014).

44. Gabriel, L., Hoff, K. J., Brůna, T., Borodovsky, M. & Stanke, M. TSEBRA: transcript selector for BRAKER. BMC Bioinformatics 22, 1–12 (2021).

45. Chen, S., Zhou, Y., Chen, Y. & Gu, J. Fastp: An ultra-fast all-in-one FASTQ preprocessor. Bioinformatics 34, i884–i890 (2018).

46. Kim, D., Langmead, B. & Salzberg, S. L. HISAT: a fast spliced aligner with low memory requirements. Nat Methods 12, 357–360 (2015).

47. Brůna, T., Gabriel, L. & Hoff, K. J. Navigating Eukaryotic Genome Annotation Pipelines: A Route Map to BRAKER, Galba, and TSEBRA. (2024).

48. Buchfink, B., Xie, C. & Huson, D. H. Fast and sensitive protein alignment using DIAMOND. Nat Methods 12, 59–60 (2014).

49. Mistry, J. et al. Pfam: The protein families database in 2021. Nucleic Acids Res 49, D412–D419 (2021).

50. Letunic, I. & Bork, P. 20 years of the SMART protein domain annotation resource. Nucleic Acids Res 46, D493–D496 (2018).

51. Haft, D. H. et al. TIGRFAMs: A protein family resource for the functional identification of proteins. Nucleic Acids Res 29, 41–43 (2001).

52. Akiva, E. et al. The Structure-Function Linkage Database. Nucleic Acids Res 42, 521– 530 (2014).

53. Pedruzzi, I. et al. HAMAP in 2015: Updates to the protein family classification and annotation system. Nucleic Acids Res 43, D1064–D1070 (2015).

54. Wang, J. et al. The conserved domain database in 2023. Nucleic Acids Res 51, D384– D388 (2023).

55. Pandurangan, A. P., Stahlhacke, J., Oates, M. E., Smithers, B. & Gough, J. The SUPERFAMILY 2.0 database: A significant proteome update and a new webserver. Nucleic Acids Res 47, D490–D494 (2019).

56. Attwood, T. K. et al. The PRINTS database: A fine-grained protein sequence annotation and analysis resource-its status in 2012. Database 2012, 1–9 (2012).

57. Mi, H., Muruganujan, A., Ebert, D., Huang, X. & Thomas, P. D. PANTHER version 14: More genomes, a new PANTHER GO-slim and improvements in enrichment analysis tools. Nucleic Acids Res 47, D419–D426 (2019).

58. Lewis, T. E. et al. Gene3D: Extensive prediction of globular domains in proteins. Nucleic Acids Res 46, D435–D439 (2018).

59. Jones, P. et al. InterProScan 5: Genome-scale protein function classification. Bioinformatics 30, 1236–1240 (2014).

60. Buenrostro, J. D., Wu, B., Chang, H. Y. & Greenleaf, W. J. ATAC-seq: A Method for Assaying Chromatin Accessibility Genome-Wide. Curr Protoc Mol Biol 109, (2015).

61. Vasimuddin, Md., Misra, S., Li, H. & Aluru, S. Efficient Architecture-Aware Acceleration of BWA-MEM for Multicore Systems. in 2019 IEEE International Parallel and Distributed Processing Symposium (IPDPS) 314–324 (IEEE, 2019). doi:10.1109/IPDPS.2019.00041.

62. Breese, M. R. & Liu, Y. NGSUtils: A software suite for analyzing and manipulating next-generation sequencing datasets. Bioinformatics 29, 494–496 (2013).

63. van der Auwera, G. & O’Connor, B. D. Genomics in the Cloud: Using Docker, GATK, and WDL in Terra. (O’Reilly Media, Incorporated, 2020).

64. Ramírez, F. et al. deepTools2: a next generation web server for deep-sequencing data analysis. Nucleic Acids Res 44, W160–W165 (2016).

65. Krueger, F. Trim Galore. https://github.com/FelixKrueger/TrimGalore (2020).

66. Rosenkranz, D. & Zischler, H. proTRAC - a software for probabilistic piRNA cluster detection, visualization and analysis. BMC Bioinformatics 13, 5 (2012).

67. Hao, Z. et al. RIdeogram: Drawing SVG graphics to visualize and map genome-wide data on the idiograms. PeerJ Comput Sci 6, 1–11 (2020).

68. Krassowski Michał. ComplexUpset.

69. Lee, J. NCBI Sequence Read Archive. https://identifiers.org/ncbi/insdc.sra:DRP012461 (2025).

70. Lee, J. NCBI Sequence Read Archive. https://identifiers.org/ncbi/insdc.sra:DRP009926 https://identifiers.org/ncbi/insdc.sra:DRP009926 (2023).

71. Lee, J. The genome annotation files of the latest genome assembly of Indian eri silkmoth (Samia ricini) including gene models, its functional annotation result, and piRNA cluster maps. FigShare https://doi.org/10.6084/m9.figshare.27268212 (2025) doi:10.6084/m9.figshare.27268212.

72. Manni, M., Berkeley, M. R., Seppey, M., Simão, F. A. & Zdobnov, E. M. BUSCO Update: Novel and Streamlined Workflows along with Broader and Deeper Phylogenetic Coverage for Scoring of Eukaryotic, Prokaryotic, and Viral Genomes. Mol Biol Evol 38, 4647–4654 (2021).

